# Stools and stool-derived extracellular vesicles from patients with Parkinson’s disease contain alpha-synuclein species with seeding capacity

**DOI:** 10.64898/2026.03.12.709633

**Authors:** Livia Civitelli, Poppy Stafford-Dorlandt, Kristijan D. Jovanoski, Afiea Begum, Selene Seoyun Lee, Elizabeth R. Dellar, Tuomas Mertsalmi, Veera Kainulainen, Perttu Arkkila, Reeta Levo, Rebekka Ortiz, Valtteri Kaasinen, Filip Scheperjans, Laura Parkkinen

**Affiliations:** University of Oxford, Nuffield Department of Clinical Neuroscience, University of Oxford, Oxford, United Kingdom; Department of Neurology, Helsinki University Hospital, and Clinicum, University of Helsinki, Helsinki, Finland; Department of Gastroenterology, Helsinki University Hospital, Helsinki, Finland; Tampere University Hospital and University of Tampere, Tampere, Finland; Clinical Neurosciences, University of Turku and Neurocenter, Turku University Hospital, Turku, Finland

**Keywords:** Parkinson’s disease, alpha-synuclein, extracellular vesicles, stools, seed amplification assay

## Abstract

**Background:** Parkinson’s disease (PD) is a neurodegenerative disorder for which there is currently no cure or reliable biomarker for early detection or for evaluating the effectiveness of potential treatments. PD pathology is driven by misfolding and subsequent accumulation of alpha-synuclein (αSyn) protein into pathological aggregates within neurons and glial cells. Seed amplification assay (SAA) is a highly sensitive and specific diagnostic tool developed to detect pathological αSyn species in the cerebrospinal fluid (CSF) of PD patients. However, αSyn aggregates are present in multiple tissues and biosamples, including stools. In this study, we aimed to investigate the potential diagnostic value of SAA using stool samples from PD patients and healthy controls (HC).

**Methods:** Stool samples from PD patients (n=45) and healthy controls (n=35) were analysed for the presence of αSyn species using slot blot assays with a panel of six αSyn antibodies, and ELISA assays. Samples were subjected to SAA, and the end-point products (SAA EP) were characterised using transmission electron microscopy (TEM). Extracellular vesicles (EVs) were isolated from the subset of samples (n=5 per group) using size exclusion chromatography and characterized by TEM. The seeding activity of isolated EVs was evaluated using SAA, followed by TEM analysis of SAA EP.

**Results:** Protein extracts from both PD and HC stool samples revealed pathological αSyn species in the slot blot assay using the phosphorylated αSyn antibody, pS129 and conformation-specific antibodies, MJFR-14 and 5G4. ELISA showed significantly elevated total αSyn levels in PD samples compared to HC, although no differences in aggregated αSyn levels were detected. In stool protein extracts, SAA demonstrated 55.6% sensitivity and 60% specificity. When applied to stool-derived EVs from PD patients and controls, sensitivity increased to 100%, while specificity remained at 60%. Notably, SAA applied to stool-derived EVs pre-incubated with recombinant monomeric αSyn achieved 100% sensitivity and 100% specificity.

**Conclusion:** These findings suggest that SAA applied to EVs isolated from stool samples, particularly after pre-incubation with recombinant monomeric αSyn, may serve as a valuable, non-invasive screening tool for the diagnosis of PD.

## Introduction

Parkinson’s disease (PD) is a neurodegenerative disorder characterised by the misfolding and accumulation of alpha-synuclein (αSyn), which forms intracellular neuronal aggregates known as Lewy bodies (LBs) and Lewy neurites (LNs) [1, 2]. The primary cause driving PD pathology remains unknown, and no intervention has yet been shown to delay disease progression. A major challenge in drug development is that PD is diagnosed only after motor symptoms, such as tremor and bradykinesia, appear [3]. By this stage, approximately 50% of dopaminergic neurons in the substantia nigra pars compacta (SNpc) are lost [4]. Robust diagnostic biomarkers could enable detection of prodromal stages of PD, facilitating earlier detection [5, 6].

αSyn Seed Amplification Assay (αSyn SAA) is an ultrasensitive aggregation-based method adapted from the prion field to detect αSyn in the CSF of patients with PD and other synucleinopathies [7]. αSyn SAA exploits the self-propagating nature of misfolded αSyn aggregates (seeds) to amplify them *in vitro*. Pathological αSyn present in a biofluid seeds aggregation of recombinant *α*Syn, which acts as substrate, through repeated elongation-fragmentation cycles [8]. Aggregation can be monitored in real-time using the amyloid dye, Thioflavin T (ThT), which emits a fluorescent signal upon amyloid fibril formation. Distinct kinetic parameters extracted from the fluorescence signal may correlate with clinical features or outcomes [9, 10]. Most studies using CSF in αSyn SAA for diagnostic purposes have reported sensitivities and specificities ≥ 90%, underscoring the utility of the technique [7, 8, 10–16]. Thus, αSyn SAA has recently contributed to a paradigm shift in PD diagnostics, moving the focus from purely clinical assessment toward molecular-based diagnosis [17, 18].

αSyn pathology is also observed in peripheral tissues, particularly within autonomic nerve terminals. The presence of αSyn aggregates in the enteric nervous system (ENS) has led to the hypothesis that PD may originate in the gut, where environmental factors trigger αSyn misfolding [19]. Pathology is thought to spread retrogradely to the brain via the vagus nerve, reaching key brainstem regions such as the dorsal motor nucleus of the vagus. Animal models support this gut-brain-axis hypothesis, αSyn aggregation and propagation after injection of preformed fibrils (PFFs) into the gut or exposure to the pesticide rotenone [20–24]. This model may explain early gastrointestinal symptoms seen in PD, which often precede motor signs by 5-10 years [25]. Epidemiological studies indicate that individuals with gastrointestinal (GI) tract disorders, such as constipation or inflammatory bowel disease, may have an increased risk of developing PD [26, 27]. Analysis of GI tissue from both PD patients and healthy controls, collected up to eight years before motor symptom onset, revealed pathological αSyn accumulation throughout GI tract [28, 29].

Extracellular vesicles (EVs) are proposed as a mechanism for systematic spread of PD pathology [30]. EVs are released by all cell types into blood circulation and contain cargo that reflects donor cell’s content. Pathological αSyn has been detected in EVs from cultured neurons [31, 32] and *in vivo* [33, 34]. Neuronally-derived EVs, immunocaptured from serum using anti-L1CAM antibodies (presumed neuronally derived), show elevated αSyn levels in PD patients and at-risk individuals, such as patients with isolated rapid eye movement sleep behaviour disorder (iRBD), a strongest predictor of phenoconversion to synucleinopathy [35]. Moreover, L1CAM-captured EVs from blood (either serum or plasma) exhibit seeding activity in αSyn SAA with high sensitivity and specificity [36, 37].

In this study, we aimed to determine whether αSyn-SAA applied to stool can detect PD-associated seeding activity, and whether stool-derived EVs harbor seeding-competent αSyn. We assessed seeding activity by αSyn-SAA in stool extracts and in isolated EV fractions, alongside biochemical characterization of stool-derived αSyn species.

## Materials and methods

### Sample collection

This study included a total of 80 stool samples that were collected at home into empty containers and were stored at the patients’ home freezer at −20C and transported to the study sites for the study visit. During the visits, samples were transferred to −80C for storage. Samples were later sent on dry ice to the lab for analysis: 45 from patients with PD (10 from the Helsinki PD cohort [38] and 35 from a PD faecal microbiota transplantation (FMT) study) and 35 from healthy controls (HC) (10 from the Helsinki PD cohort and 25 from a study on Irritable Bowel Syndrome (IBS) study) with no signs of parkinsonism or potential premotor symptoms [39, 40]. Parkinsonian symptoms were assessed using the International Parkinson and Movement Disorder Society - Unified Parkinson’s Disease Rating Scale (MDS-UPDRS III) in the FMT study, the UPDRS III in the Helsinki PD cohort and the modified Hoehn & Yahr scale (H&Y)[41]. The PD Helsinki cohort UPDRS values were converted to MDS-UPDRS by adding 7 points, based on [42]. Demographic information for all subjects is provided in Table 1. The study was approved by the Ethics Committee of the Hospital District of Helsinki and Uusimaa, and all participants provided written informed consent.

**Table 1:**
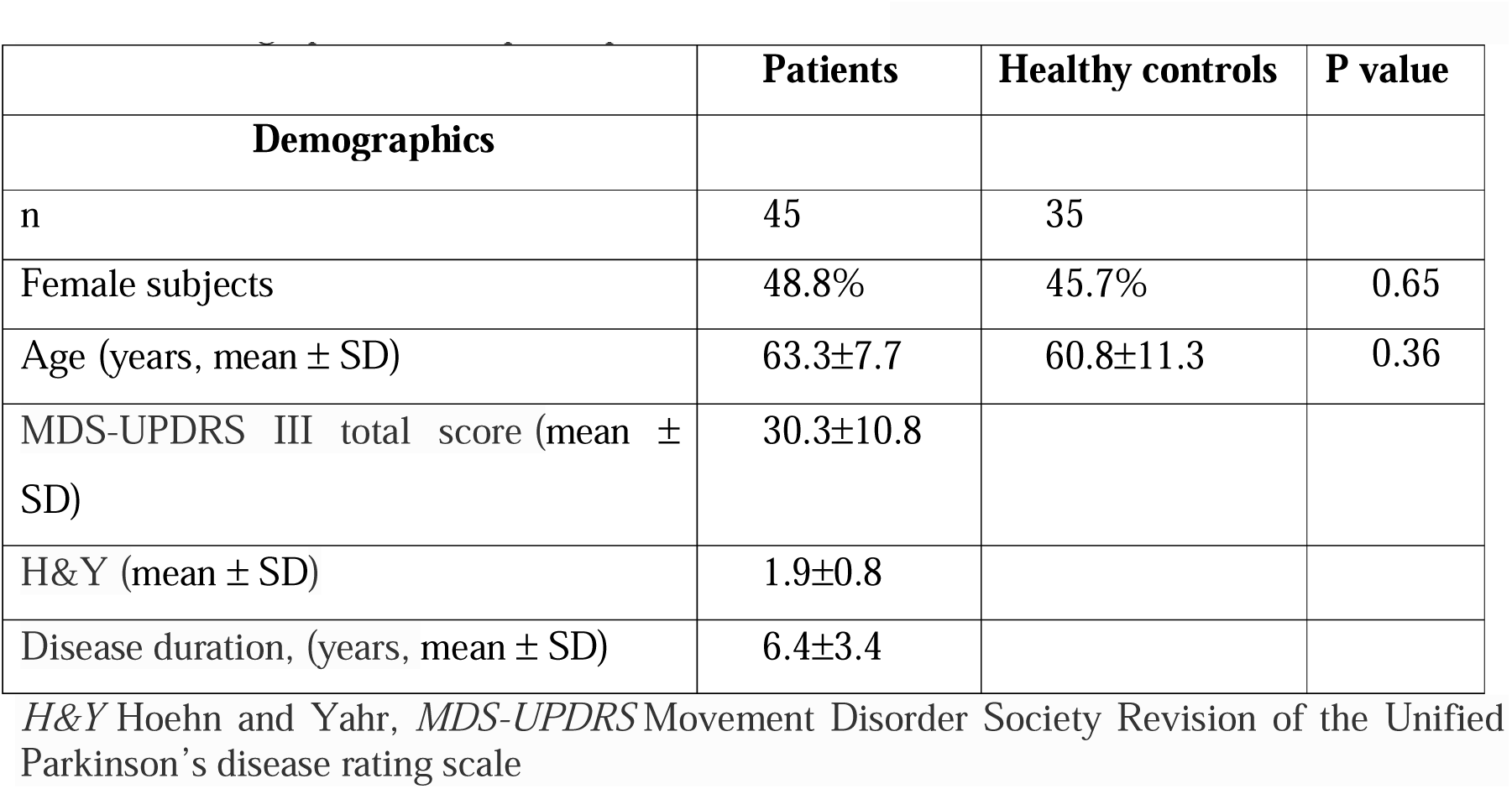
Demographic data of participants.

|  | Patients | Healthy controls | P value |
| --- | --- | --- | --- |
| **Demographics** |  |  |  |
| n | 45 | 35 |  |
| Female subjects | 48.8% | 45.7% | 0.65 |
| Age (years, mean ± SD) | 63.3±7.7 | 60.8±11.3 | 0.36 |
| MDS-UPDRS III total score (mean ± SD) | 30.3±10.8 | | |
| H&Y (mean ± SD) | 1.9±0.8 | | |
| Disease duration, (years, mean ± SD) | 6.4±3.4 | | |

*H&Y* Hoehn and Yahr, *MDS-UPDRS* Movement Disorder Society Revision of the Unified Parkinson's disease rating scale

### Protein extraction from stools

The protein extraction protocol was adapted from [43]. Briefly, frozen stool samples were resuspended in PBS containing 0.1% Tween 20 (100-300 mg/ml, 10% w/v) and vortexed at maximum speed (1400 rpm) for 20 minutes. Samples were then centrifuged at 14,000 g for 10 min at 4°C. The resulting clear supernatants were aliquoted and stored at −80°C for further use.

### Slot blot assay

Two µg of protein, prepared under non-denaturing conditions (without β-Mercaptoethanol), were immobilized onto a 0.2 μm nitrocellulose membrane (Amersham) by filtration using a slot blot apparatus (GE Healthcare) for 15 minutes. Membranes were then washed twice with PBS (2 x 10 minutes) and blocked with 5% skimmed milk in 0.1 % TBS-Tween for 1 hour at room temperature (RT). The following primary antibodies were used: anti-alpha-synuclein (phospho-Ser129), anti-p*α*Syn, 1:2000, Abcam), MJFR-14-6-4-2 (1:5000, Abcam), anti-aggregated α-synuclein, clone 5G4 (1:3000, Merck Millipore), LB509 (1:2000, Abcam), SynO4 (1:5000, a kind gift from Prof. Omar El Agnaf) and HL1-20 (1:3,000, a kind gift from Prof. Hilal Lashuel). Antibodies were diluted in 5% BSA, and membranes were incubated overnight at 4C. Membranes were subsequently washed three times with 0.1 % TBS-Tween (3 x 10 minutes) and incubated with HPR-conjugated goat anti-rabbit or anti-mouse secondary antibodies (dilution 1:5,000). Recombinant monomeric αSyn (0.1 µg) and brain homogenate (BH, 2 µg) were used as controls to verify antibody specificity. The signal from recombinant monomeric αSyn was used to assess background immunoreactivity, which was subtracted from the tested samples. Samples showing a positive signal after background subtraction were considered reactive to the antibody and assigned a binary category (0 = no reaction, 1 = reaction) for further analysis. Membranes were visualized using the ECL Western blot detection kit (Amersham). Cross-reactivity of stool samples with secondary antibodies was tested for both goat anti-mouse and anti-rabbit, confirming the absence of non-specific binding (data not shown).

### Coomassie Blue Staining

One μg of recombinant monomeric αSyn and two μg of brain homogenates were loaded on to 12% SDS-PAGE gel. Samples were prepared with Laemmli sample buffer without β-mercaptoethanol. Proteins were visualized using a Coomassie Blue staining kit according to the manufacturer’s instructions (Imperial Protein Stain, ThermoScientific). Coomassie staining of the recombinant monomeric *α*Syn, used as a control for each slot blot, confirmed that the monomeric (15 kDa) *α*Syn was predominantly present, with a minor fraction of dimeric (35 kDa) *α*Syn also detected (Supplementary Figure 1).

### Immunoblotting

Immunoblotting was carried out using a tris-glycine gel system for separation (Novex, Invitrogen). EVs were lysed directly in 2x SDS sample buffer without NuPAGE dithiothreitol reducing agent (Invitrogen), and frozen at −80 °C. Before loading, samples were heated to 65C for 12 min. Samples were separated on WedgeWell 8–16% tris-glycine mini gels in tris-glycine SDS running buffer alongside Spectra multicolour broad range protein ladder (Fisher Scientific) for 90 min at 150 V. Proteins were dry transferred to 0.2 μm PVDF membranes using an iBlot2 system (20 V for 1 min, 23 V for 4 min, 25 V for 1 min). After blocking in 5% milk in 0.1%-TBS-Tween for 1 h, membranes were incubated overnight with primary antibodies; CD63 (Thermofisher 10628D) or ApoB (Abcam 139401) at 1:1000 in 1% milk/0.1% TBS-Tween. After washing three times in 0.1%-TBS-Tween, membranes were incubated with rabbit or mouse horseradish peroxidase-labelled secondary antibodies (VWR International) at 1:10,000 for 1 h at room temperature. Detection was carried out with ECL Prime (Cytiva) on ChemiDoc MP (BioRad).

### Total protein measurement and ELISA assays

The total protein concentration of the stool samples was measured using the bicinchoninic acid (BCA) assay (PierceTM Rapid Gold BCA Protein Assay Kit, Thermo Fisher) following the manufacturer’s instructions. For each assay, 50 μl of stool sample was tested in duplicate, and absorbance was measured at 480 nm using a FLUOstar Omega microplate reader (BMG Labtech). Levels of total αSyn and aggregated αSyn were quantified using ELISA (LEGEND MAX human αSyn ELISA kit and Human αSyn PATHO ELISA, Roboscreen Diagnostics), respectively. The detection ranges of the total αSyn and aggregated αSyn ELISA kits were 10.2–650 and 5-250 pg/mL, respectively. The minimum detectable concentration (sensitivity) for the total αSyn ELISA kit was 1.80 pg/mL, while the sensitivity for the PATHO ELISA was determined according to positive controls provided in the batch-specific certificate.

### αSyn seeding amplification assay (SAA)

Stool samples were diluted 1:1,000 in PBS, and 2 μl was added to 98 μl of the reaction mixture (RB) consisting of 100 mM PIPES (pH 7), 150 mM NaCl, 0.05 mg/ml of recombinant monomeric αSyn (rPeptide, lot# r060622AS) and 10 μM Thioflavin-T (ThT), to a final volume of 100 μl. Three 1 mm zirconia/silica beads (Thistle Scientific) were pre-loaded per well into black 96-well plates with clear bottoms (Nalgene Nunc International). Before use, recombinant αSyn was filtered through a 100 kDa molecular weight cut-off (MWCO) filter (Ball Corporation) and centrifuged for 10 min at 3,300 x g. For comparison a subset of samples was run under alternative buffer conditions: 40mM PB (pH 8.2), 160 mM NaCl, 0.00125% SDS, 0.1 mg/ml of recombinant αSyn, and 10 μM ThT, with wells pre-loaded with four beads (DAIHAN Scientific). Plates were sealed with a plate sealing film (Thermo Scientific) and incubated in a FLUOstar Omega plate reader (BMG Labtech) at 42°C with shaking cycles of 1 min on (400 rpm, double orbital) and 1 min off for approximately 80-90 hrs. ThT fluorescence was measured every hour (450 ± 10 nm excitation; 480 ±10 nm emission; bottom read). The ThT fluorescence threshold for each run was calculated as the average fluorescence of all samples between cycles at 2 and 6 hours plus 10 standard deviations (SD). Each stool sample was run in quadruplicate and considered positive if at least two out of four replicates crossed the threshold.

EVs were diluted 1:1,000 in PBS, and 2 μl was added to 98 μl of the reaction mixture as above. EV samples were run in triplicate using PIPES buffer as above. A sample was considered positive if at least two out of three replicates crossed the threshold.

Kinetic parameters from each reaction were extrapolated, including: *i)* T_50_ (time to reach 50 % of the maximum fluorescence); *ii)* TTT (time needed to cross the threshold, > 10 SD above minimum fluorescence); *iii)* AUC (area under the curve over a given interval t_1_-t_2_); *iv)* F_max_ (maximum ThT fluorescence at plateau) and *v)* V_max_ (maximum slope of amplification defined as the maximum increase in relative fluorescence over time). SAA end-point (SAA EP) products were ultracentrifuged at 100,000 g for 1 hr at 4°C, collected, and stored at −80C for follow-up experiments.

### Pre-incubation of EVs with recombinant monomeric αSyn

Two μl of intact EVs, already diluted 1:1000 in PBS, were mixed in 1.5 ml Eppendorf tubes with 5 μl recombinant monomeric αSyn at 0.05 mg/ml and incubated for 5, 10, 20, 45 and 60 minutes (t=5, t=10, t=20, t=45, t=60) at 37°C in a thermoshaker (total V= 7 μl). At the end of the incubation period, the mixture was added to 93 μl of RB, and αSyn SAA was conducted as stated above. EVs were also incubated with 5 μl of recombinant monomeric αSyn immediately before adding the mixture to RB (t=0). For negative stain analysis, 20 μl of mixture at time points t=0 and t=60 was collected before running SAA.

### Transmission electron microscopy and immunolabeling electron microscopy

Ten μl of SAA EP were applied to glow-discharged carbon grids (Agar Scientific, 300 mesh) and incubated for 2 min. The grids were washed with water for 10 sec and negatively stained with 2% uranyl acetate for 10 seconds. Stained samples were imaged on a JEOL 1400 Flash microscope operated at 120 kV. For immunogold labeling, SAA EP were applied to glow-discharged carbon grids for 30 min and then blocked with blocking buffer (0.2% fish gelatin in PBS). The grids were then incubated with anti-phosphorylated αSyn (EP1536Y, Abcam, 1:20 dilution) and anti-alpha-synuclein aggregate antibody (MJFR-14-6-4-2, Abcam, 1:50 dilution) antibodies. After washing with blocking buffer, the samples were incubated with goat anti-rabbit IgG conjugated to 10 nm gold (Ab27234, Abcam) diluted 1:10 in blocking buffer. The grids were then fixed with 0.1% glutaraldehyde in PBS and stained with 2% uranyl acetate for 30 sec. For the negative staining of EVs, pioloform film on 3 mm, 300-mesh grids without carbon layer were used (AgarScientific).

### Size Exclusion Chromatography (SEC) for isolation of EVs

Stool samples were passed through a 0.22 μm polyetherosulfone filter (Merck Millipore) and adjusted to a total volume of 2 ml with PBS (without MgCl_2_ or CaCl_2_). Size-exclusion chromatography was performed with an ÄKTA GO chromatography system with a 26 ml column packed with Sepharose 4 Fast Flow resin (GE Life Sciences; exclusion limit ∼3 × 10⁷ Da). The sample was injected and eluted in PBS at a flow rate of 1 ml/min, and EVs were collected from the 7.5 – 10.5 ml elution volume. Three ml EVs sample was further concentrated as needed using 4 ml Amicon 10 kDa MWCO centrifugal filters at 3,500 g and stored at −80 °C in aliquots.

### Nanoparticle Tracking Analysis (NTA)

Particle counting was performed using a Zetaview nanoparticle tracking instrument (Particle Metrix), calibrated with 100 nm silica microspheres (Polysciences Inc). Samples were diluted in calcium and magnesium-free PBS to achieve 100-200 particles per frame. Data were acquired at RT with the following settings: sensitivity 80, shutter 100, frame rate 30, cycles 2, minimum brightness 20, maximum size 1000, minimum size 10, and minimum trace length 15. Measurements were taken at all 11 positions and analysed using ZetaView software (version 8.05.12 SP2).

### Statistical analysis

For all statistical analyses, *p* ≤ 0.05 was considered significant. GraphPad Prism (version 10.6.1, GraphPad Software) was used to perform the following analyses: chi-squared test categorical variables in Table 1, Mann–Whitney U test for ELISA comparisons (Figures 1J–K) and quantitative comparisons of the kinetic parameters for samples that amplified during SAA (Figures 2D, 3C, 4B, 4D), Spearman and Pearson correlation coefficients between SAA T_50_ and clinical characteristics (Figures 2E–H), and two-way ANOVA followed by multiple comparisons (Tukey’s HSD) for pre-incubation timepoints (Figure 4C).

**Figure 1.**
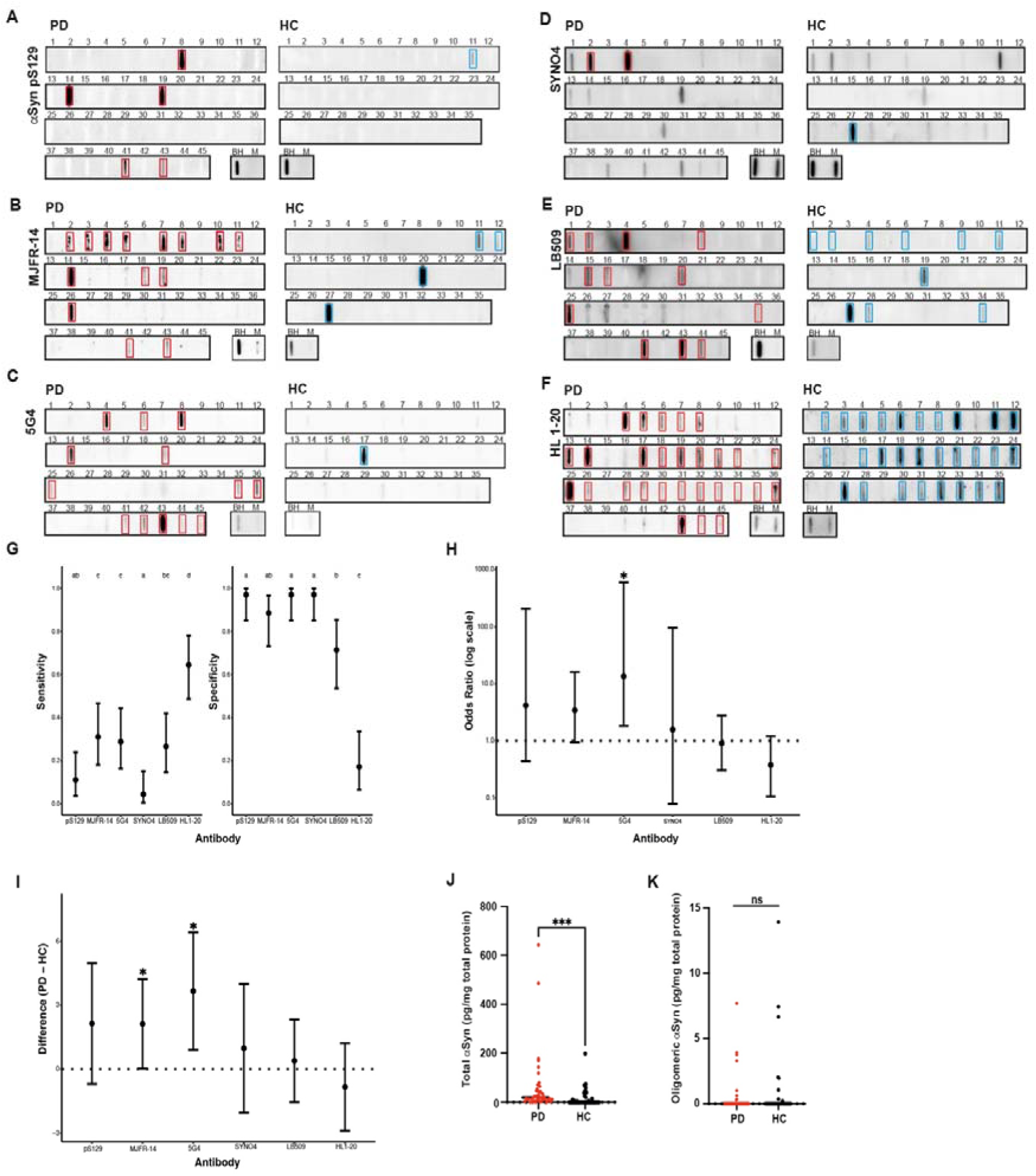
The antibody 5G4 discriminates between stool extracted from PD patients and healthy controls. Slot blots showing the immunoreactivity of stools from PD patients and healthy controls under native conditions using the following antibodies (A) anti-p*α*Syn detecting phosphorylation at Ser129, (B) MJFR-14 detecting filamentous αSyn aggregates, (C) 5G4 and (D) SYNO4 detecting conformation-specific αSyn aggregates, (E) LB509 against the C-terminus of *α*Syn, and (F) HL1-20 against the N-terminus. (BH = brain homogenate; M = recombinant monomeric αSyn). Statistically significant signals in the slot blot are indicated by red squares (PD) or light blue squares (HC). (G) Summary table showing slot blot sensitivity and specificity for each antibody (%, [95% CI]). Different letters above groups statistically significantly differences (Cochran’s Q test followed by McNemar’s pairwise comparisons; *p* < 0.05). (H) Odds ratios for each antibody’s reactivity to PD versus HC samples (estimate ± 95% CI). Asterisks indicate statistical significance (Fisher’s exact test; *p* < 0.05). (I) GLMM pairwise contrasts comparing reactivity to PD vs HC samples for each antibody (estimate ± 95% CI). Asterisks indicate statistical significance (*p* < 0.05). Quantitative ELISA analysis of stools from PD patients and healthy controls showing levels of total (J) and aggregated (K) αSyn. Results are presented as \*\*\**p*<0.001 (Mann-Whitney U test).

**Figure 2.**
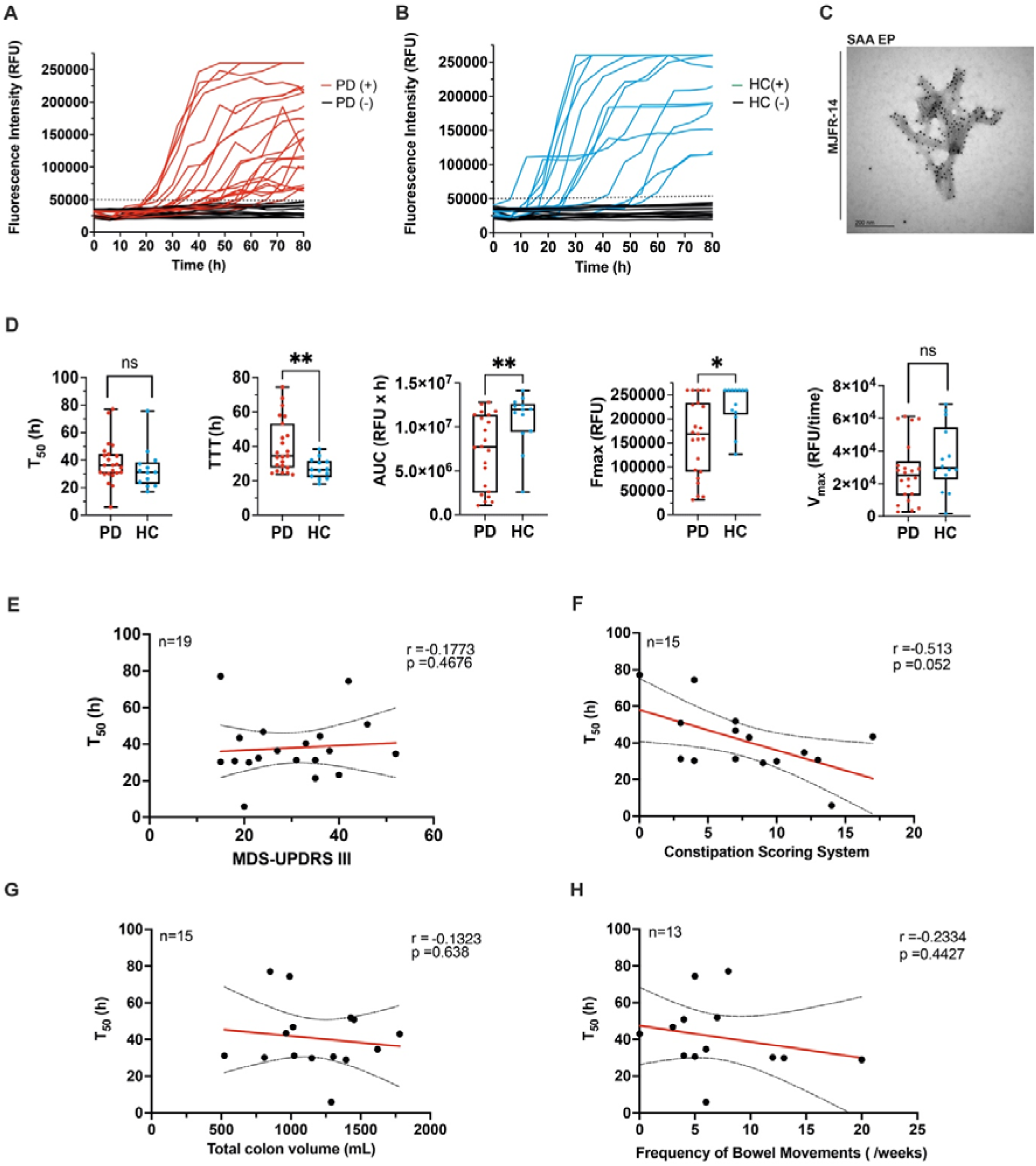
αSyn SAA seeded with stool extracts from PD patients and healthy controls. αSyn SAA was performed using stool extracts from 45 PD patients (A) and 35 healthy controls (HC) (B). Curves represent the average of four replicates per sample. C) Electron microscopy image of end-point SAA fibrils (SAA EP) from a representative PD sample, labelled with the fibril conformation-specific antibody MJFR-14 (scale bar=200 nm). D) Kinetic parameters of positive SAA reactions: T_50_, time to reach 50% of the maximum fluorescence; TTT, lag phase defined as the time when fluorescence first exceeded the threshold (50 000 RFU); AUC, area under the curve; F_max_, maximum fluorescence; V_max_, maximal rate of fluorescence increase. Each dot represents the mean of four technical replicates per sample. Statistical comparisons were performed using the Mann-Whitney U Test (\**p*<0.05, \*\**p*<0.01, ns = not significant). E-H) Correlation of T_50_ with clinical and gut-related variables: E) MDS-UPDRS III motor score F) Wexner constipation score (FMT cohort). G) total colon volume (FMT cohort), and H) bowel movement frequency score (FMT cohort). Each dot represents an individual patient sample. Correlations were assessed using Spearman’s rank correlation. For illustration, scatter plots also show Pearson correlation coefficient (r), linear regression lines (red), 95% confidence intervals (dotted lines), and corresponding *p*-values.

R (version 4.5.2) was used to calculate sensitivity and specificity (epiR package, version 2.0.88) with 95% confidence intervals using the Clopper-Pearson exact method for slot blot (Figures 1G) and SAA (Figure 4F), Cochran’s Q test (RVAideMemoire package, version 0.9.83.12) followed by McNemar’s pairwise comparisons (rcompanion package version 2.5.0) for antibody immunoreactivity (Figure 1G), Fisher’s exact test for PD versus HC responses in slot blot (Figure 1H) and SAA (Figure 4G), and а generalized linear mixed-effects model (GLMM) followed by pairwise comparisons for slot blot (Figure 1I).

Тhe GLMM was fitted using the glmer() function from the lme4 R package (version 1.1.37) with the formula "slotblot ∼ antibody * state + cohort + sex + age + (1 | sampleID)”. The response variable “slotblot” was binary (0 = no reaction, 1 = reaction). Fixed effects included “antibody” (pS129, MJFR-14, 5G4, LB509, SYNO4, or HL1-20), disease “state” (PD or HC), their interaction (“antibody” × “state”), “cohort” (Helsinki, IBS, or FMT), “sex” (female/male), and “age” (rounded down to the nearest integer). Donor identity (“sampleID”) was modelled as a random intercept to account for repeated measurements for within donors. Model fitting used the BOBYQA optimizer with a maximum of 200,000 iterations.

GLMM fixed-effect estimates were calculated as log-odds with 95% Wald confidence intervals and exponentiated to obtain odds ratios. Post hoc pairwise contrasts (PD versus HC for each antibody) were estimated using the emmeans R package (version 2.0.0). Estimated marginal means were calculated on a log-odds scale and transformed to odds ratios with p-values for contrasts derived from Wald tests.

## Results

### Stools from Parkinson’s patients contain pathological αSyn species

Demographic and clinical information for the study participants is provided in Table 1. Protein was extracted from the stools of 45 PD patients and 35 HC and analysed by slot blot using a panel of six antibodies against αSyn (Figure 1A-F). Only five samples (IDs #8, #14, #19, #41, #43) showed immunoreactivity with pS129 (anti-p*α*Syn, Figure 1A), these same samples also reacted with MJFR-14 (Figure 1B) and 5G4 (Figure 1C), which preferentially detect pathological *α*Syn species [44, 45]. First, we determined the sensitivity and specificity of each antibody (Figure 1G). While all six antibodies had low sensitivity (all < 65%, five < 32%), three antibodies (pS129, 5G4, and SYNO4) had very high specificity (97.1%). Although HL1-20 had the highest sensitivity (64.4%), it also showed the lowest specificity (17.1%). Of the three antibodies with the highest specificity, 5G4 had the highest sensitivity (28.9%) (Figures 1G).

To assess whether 5G4 could discriminate between PD and HC samples better than the other antibodies, we performed two independent statistical analyses. Using Fisher’s exact test, only 5G4 demonstrated a statistically significant odds ratio for reactivity to PD versus HC samples (13.5, 95% CI 1.83–602, *p* = 0.0024, Figure 1H). We then fitted a generalized linear mixed model (GLMM) to the binary slot blot data, accounting for sex, age, cohort, repeated measures, and interactions between antibody and disease state (see Materials and Methods). Pairwise contrasts from the GLMM (PD minus HC reactivity) indicated statistically significant differences for 5G4 (*p* = 0.00936) and MJFR-14 (*p* = 0.0475) (Figure 1I). Together, these results suggest that 5G4, which preferentially binds to pathological *α*Syn aggregates [40], can discriminate stool samples from PD patients and healthy controls.

Next, we assessed total and aggregated Syn concentrations in stools using ELISA assays. Total αSyn levels were significantly higher in PD patients compared to HC (*p*<0.001, Figure 1J), and this remained significant after excluding two outliers (data not shown). Oligomeric αSyn levels were measured using the Patho ELISA assay, which is based on the 5G4 antibody [45] and has a sensitivity range of 5 – 250 pg/ml. However, in 35 of 45 PD samples, no αSyn aggregates were detected, and in 3 PD samples, levels were below the detection limit, thus these samples were excluded from the analysis. Among the HC control group, 27 samples showed no detectable αSyn aggregates, and two had levels below the detection limit, resulting in their exclusion. Overall, the Patho ELISA assay did not detect higher concentration of aggregated αSyn species in PD stools compared to HC *(p* = 0.539, Figure 1K).

### Stools from Parkinson’s patients show seeding activity

We next explored whether stool samples could be used in the αSyn seeding amplification assay (SAA) as an alternative to established biofluids such as CSF [10]. To optimise the assay, we compared two buffer conditions, PIPES versus phosphate buffer (PB) using stool extracts from 5 PD patients and 5 healthy controls (HC) (Supplementary Figure 2). Using PIPES, 3 of 5 PD and 2 of 5 HC samples showed amplification, with generally higher fluorescence signals, including one PD sample reaching a strong signal. In contrast, with PB buffer, 3 of 5 HC and only 1 PD sample amplified; HC samples showed faster seeding activity, but overall fluorescent was very low. Based on these results, we selected PIPES buffer for the full cohort.

Stool samples from 45 PD patients and 35 HC were diluted 1: 1,000 and run in quadruplicates alongside negative controls (recombinant αSyn alone) for 80 hours. Samples were considered positive when fluorescence crossed the threshold (dotted line). From each reaction, kinetic parameters were extracted: *i)* T_50_, time to reach 50 % of maximum fluorescence; *ii)* TTT, the time to cross the threshold (> 10 SD of the minimum fluorescence); *iii)* AUC, area under the curve between defined timepoints; *iv)* F_max_, maximum ThT fluorescence at plateau and *v)* V_max_, maximum slope of fluorescence increase over time).

Overall, 25 of 45 PD samples and 14 of 35 HC samples were positive for αSyn SAA, corresponding to a sensitivity of 55.6% (95% CI, 40.0–70.4%) and specificity of 60.0% (95% CI, 42.1–76.1%), (Figure 2A-B). To confirm the identity of amplified fibrils at the SAA end point (SAA EP), we performed immunogold-TEM using the αSyn fibril-specific antibody MJFR-14, verifying that the amplified structures were indeed αSyn (Figure 2C). Kinetic parameters did not reveal statistically significant differences in seeding activity between the diagnostic groups overall. However, when analysing only positive samples, PD and HC differed significantly in TTT, AUC and F_max_. Notably, stool samples from HC amplified faster (shorter TTT), and reached higher maximum fluorescence (F_max_), (Figure 2D, *p*<0.05 or *p*<0.01, not significant for some comparisons). Among PD samples that amplified, no correlation was observed between T_50_ and clinical severity measured by MDS-UPDRS III (Figure 2E). Similarly, in the subset of PD samples from the FMT study, gut-related variables including Wexner Constipation Scoring System, total colon volume, and bowel movement frequency did not correlate with T_50_ (Figure 2F–G). These findings were consistent when TTT was analyzed instead (data not shown).

### Extracellular vesicles are successfully isolated from stools in both PD patients and healthy controls and exhibit seeding activity

Extracellular vesicles (EVs) have recently been applied in αSyn SAA after isolation from plasma, yielding promising results [36, 37]. Building on this, we aimed to isolate EVs from stool extracts of 5 PD patients (PD13, PD21, PD36, PD41 and PD44) and 5 healthy controls (HC13, HC22, HC24, HC26 and HC27) from the FMT study, with a goal of enhancing the αSyn SAA performance previously shown in total stool homogenates. EVs were isolated using size exclusion chromatography (SEC), and the resulting SEC profile revealed a single peak between 7.5 ml and 10.5 ml (Supplementary Figure 3A), corresponding to the collected EVs as confirmed by nanoparticle tracking analysis (NTA, Supplementary Figure 3B). The morphology of the extracted EVs was validated by TEM, while immunoblotting confirmed the presence of EV-enriched marker (CD63) and the absence of the lipoprotein marker ApoB, supporting EV enrichment and minimal lipoprotein contamination. (Supplementary Figure 3C-D).

Intact EVs were used as seeds in the αSyn SAA assay at a dilution of 1: 1,000 to assess their capacity to induce αSyn aggregation. As shown in Figure 3, over an 80-hour reaction period, EVs from all 5 PD patients induced an increase in ThT signal, although only one sample (PD1) reached maximum fluorescence intensity (F_max_) (sensitivity = 100%, 95% CI 47.8–100%, Figure 3A). In contrast, EVs from healthy controls showed amplification in 2 out 5 samples (specificity = 60%, 95% CI 14.7–94.7%, Figure 3B). Notably, when only EVs were used in the αSyn SA without further processing, no amplification signal was revealed. Analysis of kinetic parameters (T_50_, F_max_ and V_max_) for the samples showing amplification revealed statistically significant differences between PD and HC (*p*<0.05 and *p*<0.01, Figure 3). TEM imaging of the SAA EP from PD1 and HC1 confirmed the presence of EVs incorporated within fully formed fibrils (Figure 3D, black arrow). Immunogold-TEM using the MJFR-14 antibody further validated that the SAA EP amplified in the presence of EVs consisted of αSyn.

**Figure 3.**
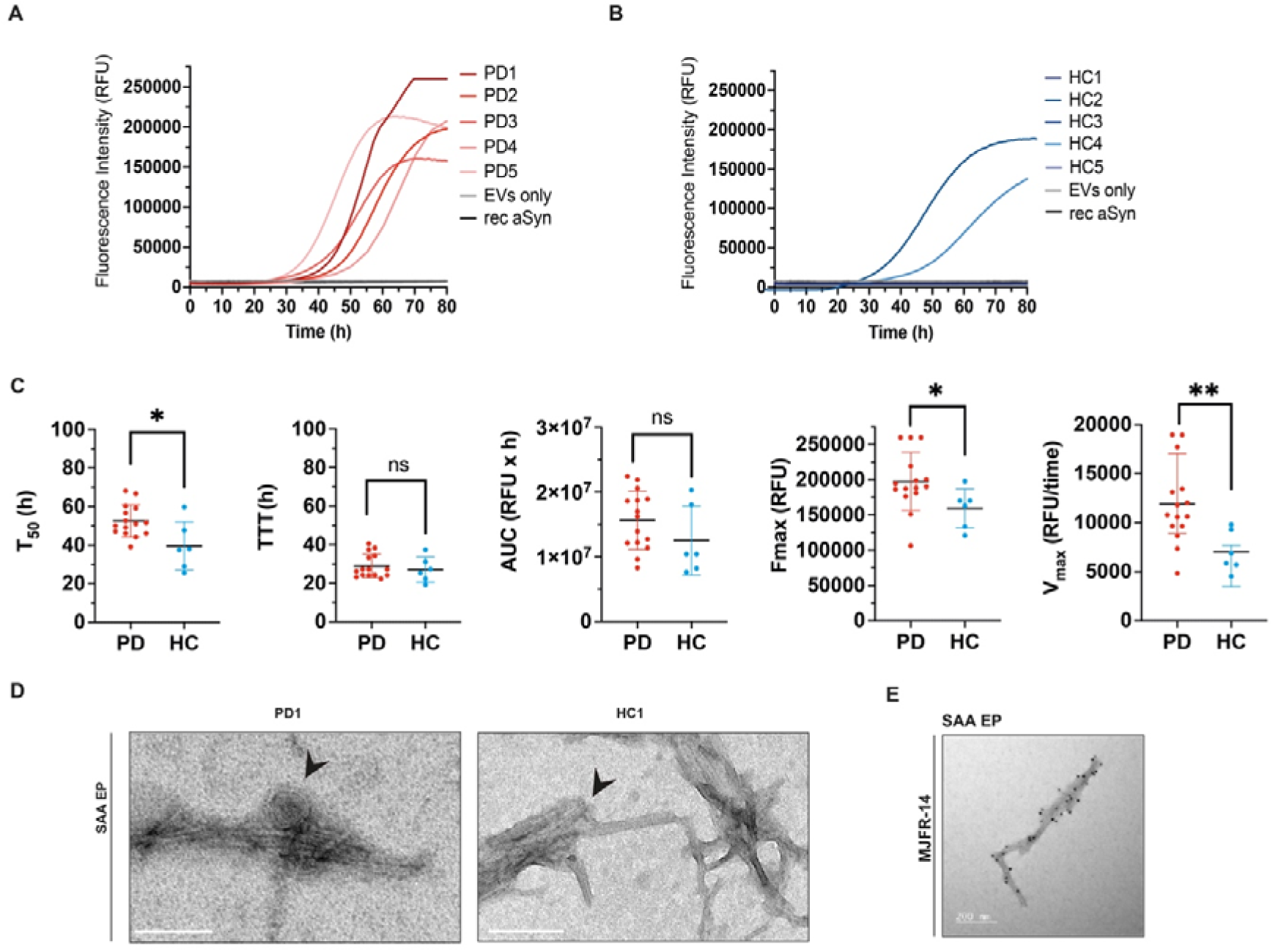
Stool-derived EVs from PD patients trigger αSyn aggregation. A) αSyn SAA performed with EVs extracted from stools of 5 PD patients and B) 5 healthy controls. C) SAA kinetic parameters derived from the assay: T_50_ = time to reach 50% of maximum fluorescence; TTT= time at which each positive reaction exceeds the threshold (30 000 RFU); Area under the curve (AUC); F_max_ = maximum ThT fluorescence at plateau; V_max_ = maximum slope of the amplification per unit of time. Each dot represents a single measurement from an individual sample within each group (Mann-Whitney U Test, \**p*<0.05, \*\**p*<0.01, ns = not significant). Each point represents an individual technical replicate (four replicates per sample). D) Transmission electron microscopy image of SAA end-point fibrils (SAA EP) from samples PD1 (left) and HC1 (right). Scale bar=400 nm. Black arrows indicate EVs incorporated in SAA EP fibrils. E) Transmission electron microscopy image of SAA EP fibrils from one representative PD sample labelled with the fibril conformation-specific antibody MJFR-14. Scale bar=200nm.

### Extracellular vesicles isolated from PD patients induce nucleation of recombinant αSyn in pre-incubation experiments

To gain insight into the stage at which EVs affect αSyn aggregation, stool-isolated EVs from PD were pre-incubated at 37°C under quiescent conditions over a time course, followed by αSyn SAA. EVs were pre-mixed with recombinant monomeric αSyn for t=5, t=10, t=20, t=45, and t=60 minutes, or immediately before running αSyn SAA experiment (t=0). During the initial incubation times (t=0, 5, 10, and 20 min), stool-derived EVs promoted αSyn fibril formation, but the overall ThT signal was low (F_max_<200,000 RFU). At t=45 and t=60, stool-isolated EVs triggered αSyn fibril formation, with four samples reaching F_max_ (260,000 RFU) (Figure 4A, sensitivity 100%, CI 47.8%-100%). TTT showed a significant difference between 5 and 10 min versus 60 min of incubation (*p* <0.05). In contrast, F_max_ was significantly higher at t=45 versus t=0 (*p* <0.001) and t=5 (*p* <0.01), and at t=60 versus t=0, 5, 10 and 20 min (*p* <0.0001), but not versus t=45 (Figure 4B). When the same experiment was performed with EVs isolated from healthy controls (HC), EVs triggered αSyn aggregation at t=0, but at t=60 the ThT signal was completely abolished, indicating that prolonged pre-incubation of recombinant monomeric αSyn with HC-derived EVs prevented αSyn aggregation (specificity 100%, CI 47.8%-100%). The calculated TTT was statistically different between t=0 and t=60 (*p*<0.0001, Supplementary Figure 6A-B). When comparing the kinetic parameters of pre-incubation experiments using PD- and HC-derived EVs, TTT was consistently shorter for PD-derived EVs at both t=0 and t=60 (Figure 4C, *p*< 0.0001). In contrast, F_max_ was significantly higher at t=0 when HC-derived were used in pre-incubation experiments (Figure 4C, *p*<0.01; *p*<0.0001). However, at t=60 this trend was reversed, with PD-derived EVs producing a significantly higher F_max_ compared to HC (Figure 4C, *p*<0.0001). The pro-aggregating effect of PD-derived EVs at t=0 was further confirmed by TEM, which revealed the presence of short fibrils immediately after incubation of recombinant αSyn with stool-derived EVs. In contrast, a longer pre-incubation period (t=60) led to the formation of short fibrils characterised by lateral associations (Figure 4D). Raw kinetic data from the technical replicates of stool-derived EVs from PD patients are shown in Supplementary Figure 5. Finally, we compared the diagnostic performance of the three different SAA methods used in our study: stool protein extracts (Figure 2), stool-derived EVs (Figure 3), and pre-incubated (t = 60) stool-derived EVs (Figure 4). SAA sensitivity was higher for stool-derived EVs than for stool protein extracts, whereas SAA specificity improved when stool-derived EVs were also pre-incubated with recombinant monomeric αSyn for 60 minutes (Table 2). Notably, only the method using pre-incubated EVs yielded a statistically significant odds ratio for amplification in PD versus HC samples (Figure 4E), albeit with an uncertain estimate and upper bound (Fisher’s exact test, CI 2.5 to ∞, *p* = 0.0008). Overall, these SAA results suggest that incubating stool-derived EVs with recombinant monomeric αSyn for 60 minutes before SAA may provide a readout that can robustly distinguishes stool samples from PD patients and healthy controls.

**Figure 4.**
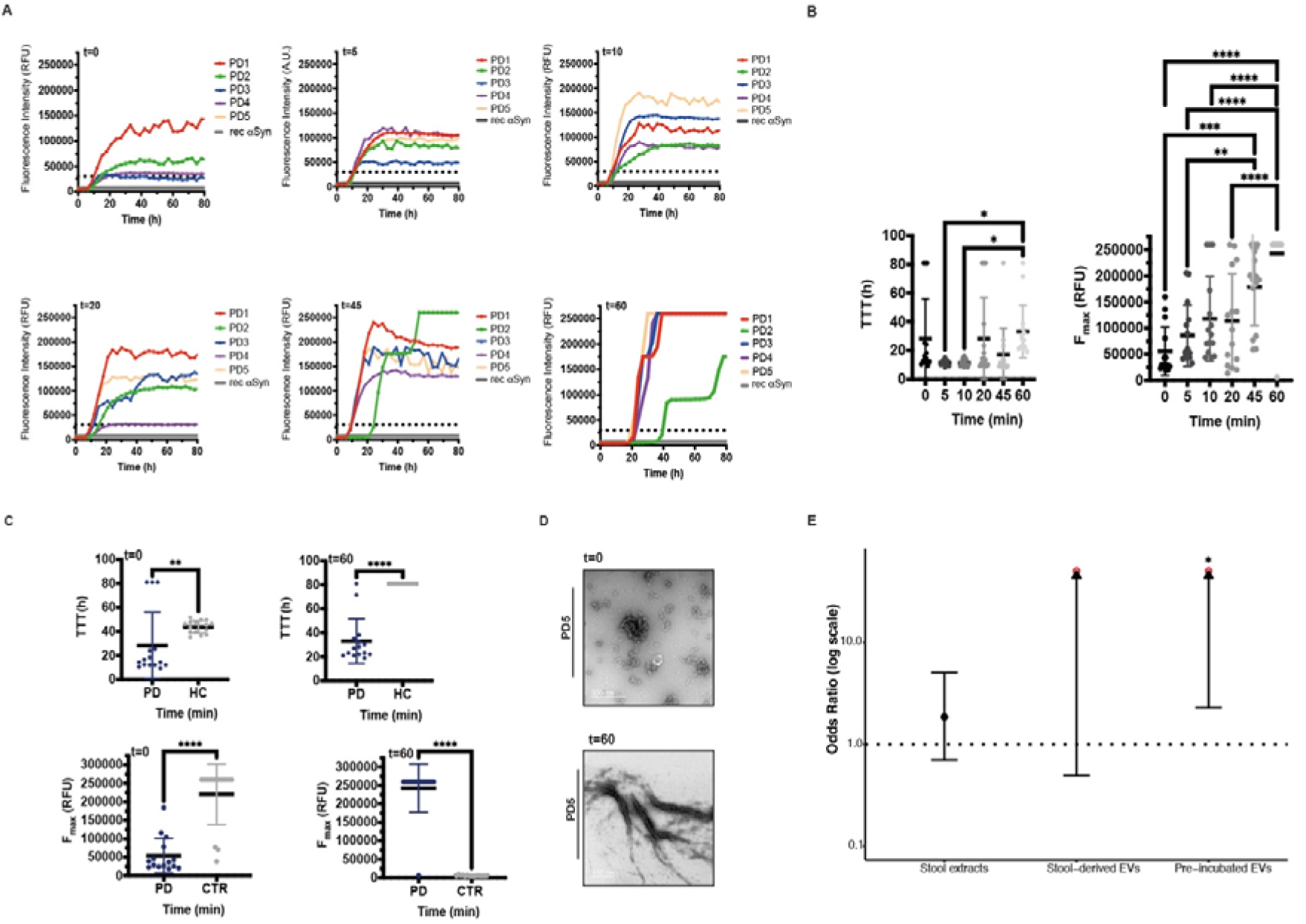
Stool-isolated EVs affect αSyn aggregation in pre-incubation experiments. A) Stool-isolated EVs from PD patients were pre-incubated with recombinant monomeric αSyn for t=0, 5, 10, 20, 45, and 60 min, followed by αSyn SAA. B) Graphical representation of TTT and F_max_ obtained from time-course pre-incubation experiments using stool-isolated EVs from PD patients (two-way ANOVA with Tukey post hoc test, \**p*<0.05, \*\**p*<0.01, \*\*\**p*<0.001, \*\*\*\**p*<0.0001). C) Graphical representation of the TTT at t=0 and t=60 for EVs isolated from PD compared with EVs isolated from HC (Mann-Whitney U Test, \*\**p*<0.01; \*\*\*\**p*<0.0001). Graphical representation of the F_max_ at t=0 and t=60 for EVs isolated from PD compared with EVs isolated from HC (Mann-Whitney U Test, \*\*\*\**p*<0.0001). Each experiment was performed in triplicate. The threshold is indicated by a dotted line and was set at 30,000 A.U. Dots represent individual technical replicates. D) Representative TEM images at t=0 (top panel) and t=60 (bottom panel). Scale bars= 500 nm and 200 nm. E) Odds ratios for each SAA method for amplification in PD versus HC samples (estimate ± 95% CI). Open red circles denote uncertain estimate (∞), and black arrowheads denote uncertain upper bounds (∞). Asterisks indicate statistical significance (Fisher’s exact test; *p* < 0.05).

**Table 2:** Summary table of sensitivity and specificity for each SAA method (%, [95% CI])

| SAA | Sensitivity (%) | Specificity (%) |
| --- | --- | --- |
| Stool extracts | 55.6 [40.0; 70.4] | 60.0 [42.1; 76.1] |
| Stool-derived EVs | 100.0 [47.8; 100.0] | 60.0 [14.7; 94.7] |
| Pre-incubated EVs | 100.0 [47.8; 100.0] | 100.0 [47.8; 100.0] |

## Discussion

In this study, stools from both Parkinson’s disease (PD) patients and healthy controls (HC) were shown to contain a spectrum of αSyn species, ranging from monomers to pathological aggregates. We further showed that αSyn seed amplification assay (αSyn SAA) can be performed directly on stool samples. Higher sensitivity was achieved when intact stool-isolated extracellular vesicles (EVs) were used as an alternative matrix. In particular, pre-incubation of intact stool-derived EVs with recombinant αSyn enabled complete separation of PD and HC samples, with 100% sensitivity and 100% specificity. Therefore, we propose that stool-derived EVs pre-incubated with recombinant αSyn might represent a potentially robust and non-invasive alternative for αSyn SAA-based detection of PD [9, 10].

The panel of antibodies used against αSyn helped us identify the diversity of αSyn species present in stools, confirming that PD patients exhibit both physiological and pathological αSyn species. The pathogenic role of phosphorylation at serine 129 (pS129) remains controversial. Some studies report that pS129 promotes αSyn aggregation [47], whereas others suggest that it inhibits αSyn aggregation with neuroprotective effects [48, 49]. Notably, more than 90% of αSyn in Lewy bodies (LBs) is phosphorylated at Ser129 [52], while only 5% of αSyn in healthy brains carries this post-translational modification. Thus, detecting pS129 in stools is particularly intriguing, as it may reflect Lewy body–like pathology in the enteric nervous system or gut-associated tissues.

MJFR-14, 5G4 and SYNO4 are all conformation-specific antibodies that preferentially recognise aggregated αSyn [44–46]. They exhibit weak immunoreactivity toward monomeric αSyn but can bind to both oligomeric and fibrillar species [47]. 5G4 preferentially recognises β-sheet–rich oligomers and fibrils, whereas SYNO4 detects more unstructured oligomers. In our slot blot experiments, 5G4 detected signals in 13 out of 45 PD patients, and statistical analysis indicated that this detection rate differs significantly from that in HCs, supporting its discriminatory potential. However, the ELISA assay using 5G4 as a capture antibody did not yield statistically significant differences. Although ELISA is generally more sensitive, this higher sensitivity may have revealed low levels of aggregated αSyn in HCs that were not detectable by slot blot, thereby reducing group differences. At the same time, most samples showed no signal or values below the detection limit, suggesting that ELISA alone may not provide a comprehensive picture of pathological αSyn species in stools.

Schaffrath and colleagues [50] reported that stools from patients with REM sleep behavior disorder (RBD) contain significantly higher levels of aggregated αSyn then stools from PD patients or HCs. They did not find significant differences between PD patients and HCs, which aligns with our findings. Their study employed a highly sensitive surface-based fluorescence intensity distribution analysis (sFIDA), a highly sensitive method combining immunoassay with confocal microscopy that can detect αSyn aggregates at femtomolar concentrations. However, both ELISA and sFIDA rely on antibodies and may therefore be limited in their ability to fully profile αSyn species. As our slot blot results suggest, comprehensive profiling may require multiple antibodies targeting different conformations.

The origin of αSyn in stools remains unclear, although αSyn is expressed in the enteric nervous system (ENS) [51]. Several gut epithelial cell types, including enteroendocrine cells (EECs) [52], and enteric glial cells express αSyn. EECs directly face the gut lumen and form synpase-like connections with αSyn-containing enteric neurons, creating a neural circuit between the gut and the ENS. Pathological αSyn could be released from the nerve terminals or accumulate intracellularly and subsequently be secreted. Previous studies examining abnormal αSyn pathology in colonic biopsies from prodromal or early PD subjects and HCs, have yielded conflicting results depending on the region analysed and the antibodies used [29, 53–55]. αSyn SAA has also been applied to intestinal tissues [13, 49], but detection has been limited by low sensitivity or reliance on post-mortem samples. More recently, application of αSyn SAA to duodenal biopsies showed improved sensitivity and specificity [56], reinforcing the diagnostic potential of the assay.

When we evaluated the seeding capacity of protein extracts, overall diagnostic accuracy was modest (55.6% sensitivity and 60% specificity). Among samples that amplified, HCs tended to amplify faster and reach higher maximum fluorescence compared to PD patients. This suggest that the stool matrix—a complex mixture of water, free proteins, bacteria, and undigested food—differs between PD and HC. In PD, such differences may reflect gut dysbiosis [57, 58] and the presence of molecules that inhibit αSyn aggregation. Lipoproteins, for example, have been shown to inhibit αSyn aggregation in CSF [59], and similar inhibitory factors may exist in stools. Further optimization of the αSyn SAA protocol is therefore warranted.

To reduce the potential matrix effects, we isolated EVs from stool samples. To our knowledge, this is the first study applying *α*Syn SAA to intact EVs extracted from stools of PD patients and HCs. Stool-derived EVs from PD samples increased the sensitivity to 100%, but EVs from HCs also promoted aggregation, resulting in only 60% specificity. This raises questions about which EV-associated biological factor(s) — particularly lipid composition —drive *α*Syn aggregation. Studies with synthetic vesicles have shown that lipid composition strongly influences *α*Syn aggregation propensity [60, 61].

We therefore tested whether isolated EVs could act as nucleation surfaces by pre-incubating them with monomeric recombinant *α*Syn. Monomeric *α*Syn is kinetically stable, but upon binding suitable surfaces, its N-terminal regions can adopt helical conformations that facilitate aggregation [61]. EVs from PD patients triggered *α*Syn aggregation in a time-dependent manner under all tested conditions. In contrast, HC-derived EVs triggered aggregation only at the baseline (t=0), but not after preincubation (t=60), suggesting a potential inhibitory effect. Depending on their chemical properties, lipid membranes can either promote or fails to enhance *α*Syn aggregation [60, 62].

We hypothesise that EVs from different sources have distinct lipid compositions that influence *α*Syn behaviour and, in PD, may favor *α*Syn aggregation. Lipids are increasingly recognised as key players in neurodegenerative diseases due to their role in membrane structure, signalling, and protein aggregation [63]. Multi-omics analyses of PD brains reveal region-specific lipid profiles [64], and dysregulated lipid metabolism may contribute to PD pathogenesis. Additionally, *α*Syn may associate with the outer EV membrane [65], where it could nucleate further aggregation.

## Conclusion

In summary, our study demonstrates that stools from PD patients contain αSyn species capable of inducing aggregation in SAA. Interestingly, seeding kinetics (T_50_ or TTT) did not correlate with clinical severity (MDS-UPDRS III) or gut-related variables, suggesting that stool αSyn seeding activity may reflect intrinsic molecular properties of αSyn or the stool matrix rather than overt disease stage or gut physiology. These findings indicate that, while stool αSyn SAA can detect aggregation-prone αSyn, matrix effects and individual variability can substantially influence kinetic readouts, highlighting the need for strategies to enhance assay sensitivity and specificity. Here, we show for the first time that intact EVs can be isolated from stools of both PD patients and healthy controls. Compared with stool protein extracts, stool-derived EVs improve SAA sensitivity, and pre-incubation with monomeric recombinant αSyn enhances specificity. Our findings suggest that EV components—such as lipids or membrane-associated αSyn—may facilitate aggregation. Collectively, these findings support the use of intact stool-derived EVs pre-incubated with recombinant αSyn as a robust and non-invasive biosample to improve αSyn detection in PD.

## Acknowledgments

We thank all the patients who donated their samples for the study. We also want to thank the Dunn School Bioimaging Facility for providing access and training for transmission electron microscopy and assisting with acquiring the imaging data.

## Funding/Support

Michael J. Fox Foundation for Parkinson’s Research

Weston Brain Institute

Parkinson’s UK

Parkinson Foundation

NIHR Oxford Biomedical Research Centre

the Research Council of Finland (295724, 310835, 342758)

the Finnish Medical Foundation

the Finnish Parkinson Foundation

the Hospital District of Helsinki, and Uusimaa (T1010NL101, TYH2020335, TYH2021336, TYH2023232)

the Emil Aaltonen Foundation (210267)

the Konung Gustav V:s och Drottning Victorias Stiftelse

the Olvi Foundation

the Yrjö Jahnsson Foundation

the Stockmann Foundation

the Sigrid Jusélius Foundation

## Author Contribution

*Concept and design:* LC, PS-D, AB, FS, LP

*Recruitment of research subjects:* TM, RL, RO, VK, FS

*Acquisition, analysis, or interpretation of data:* LC, PS-D, KDJ, AB, SLL, ERD, PA, FS, LP

*Drafting of the manuscript:* LC, PS-D, AB, KDJ, LP

*Critical review of the manuscript for important intellectual content:* LC, PS-D, KDJ, ERD, TM, VK, PA, RL, RO, VK, FS, LP

*Statistical analysis:* LC, KDJ

*Obtained funding:* FS, LP

*Supervision:* LC, LP

## Financial disclosures

FS reports consultancies (Abbvie, Axial Biotherapeutics, Orion Pharma, Teva, Adamant Health, Merz), honoraria (Abbvie, GE Healthcare, Merck, Teva, Bristol Myers Squibb, Sanofi, Biocodex, Biogen), grants (Emil Aaltonen Foundation, Konung Gustav V:s och Drottning Victorias Stiftelse, Olvi Foundation, Yrjö Jahnsson Foundation, Stockmann Foundation, Sigrid Juselius Foundation, Research Council of Finland, Hospital District of Helsinki and Uusimaa, Renishaw, Orion Pharma, Abbvie), travel expenses (Lundbeck, NordicInfucare), stock (NeuroInnovation Oy, NeuroBiome Oy, Axial Biotherapeutics).

## Notes

### Competing Interest Statement

The authors have declared no competing interest.

