## Supplementary Figures for "Stools and stool-derived extracellular vesicles from patients with Parkinson’s disease contain alpha-synuclein species with seeding capacity"

**Supplementary Figure 1. Characterization of monomeric recombinant αSyn.** SDS-PAGE followed by Coomassie blue staining revealed the presence of monomeric (15 kDa) and dimeric (35 KDa) αSyn. *Lane 1*: rain homogenate (BH); *lane 2*: monomeric recombinant αSyn (rec αSyn).

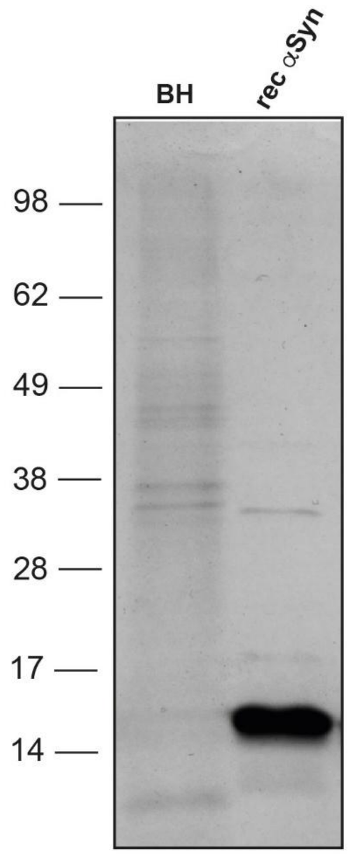

**Supplementary Figure 2. αSyn SAA reaction seeded with stool extracts using PIPES or phosphate buffer.** αSyn SAA was performed with stool extracts of 5 Parkinson´s disease patients (PD) and 5 healthy controls (HC) using either (A) 100 mM PIPES, pH 7, or (B) 40 mM phosphate buffer (PB), pH 8.2. (C-G) Raw data of αSyn SAA reactions using PIPES buffer aew shown for each sample that produced a positive ThT signal; each replicate (n=4) is shown. H) Fluorescence kinetics are shown as mean ± S.D. for each positive sample.

The threshold, indicated by a dotted line, was set at 40,000 RFU.

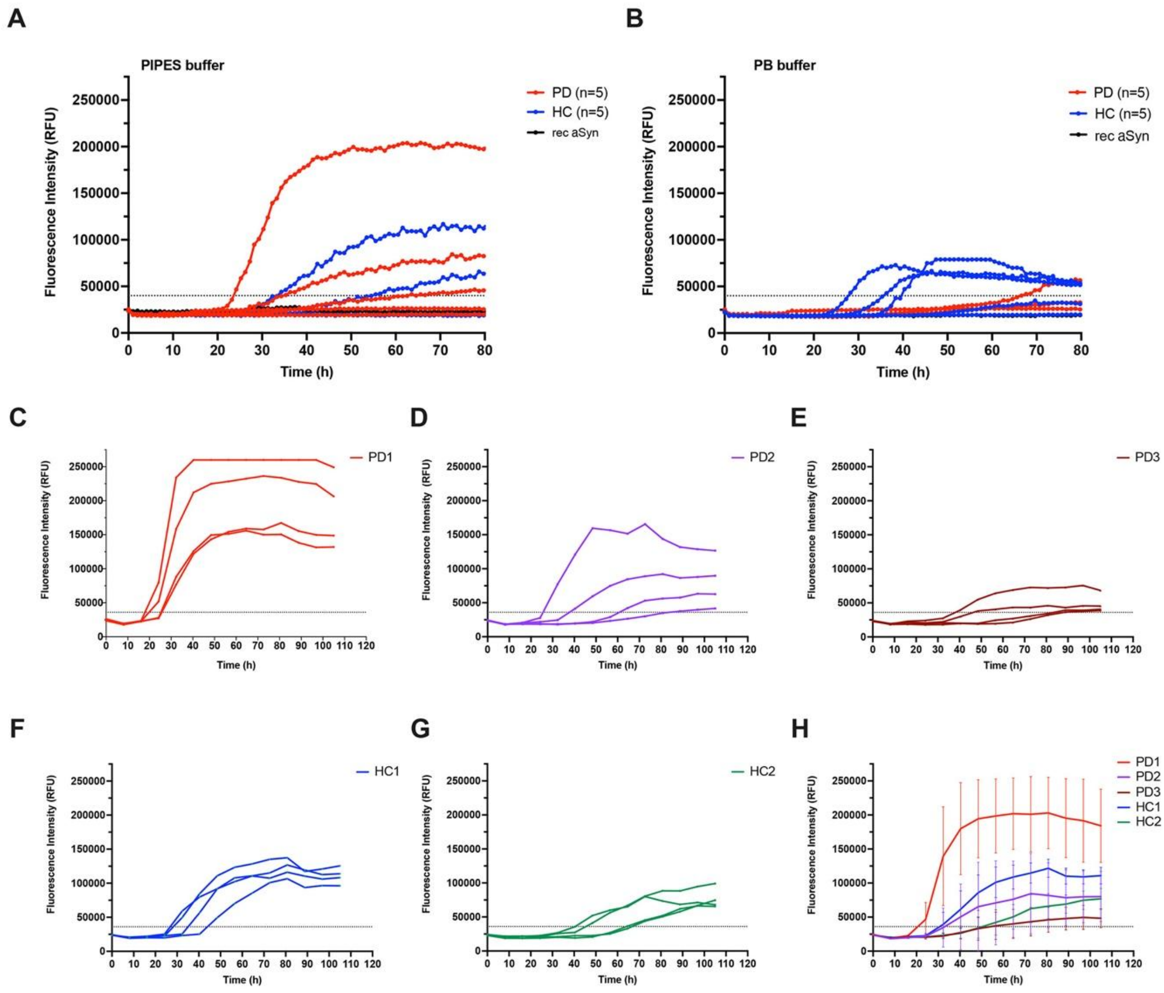

**Supplementary Figure 3: Validation of extracellular vesicle separation.** A) Size-exclusion chromatography protein trace showing the separation of extracellular vesicle-containing fractions from bulk free protein. B) Nanoparticle tracking analysis (NTA) confirming the presence of particles within expected elution volume (7.5-10.5 ml). C) Representative negative-stain transmission electron microscopy (TEM) image of pooled elution fractions (7.5-10.5 ml). Scale bar = 1000 nm. D) Representative immunoblotting image of sample PD3 probed with anti-CD63 and anti-ApoB antibodies.

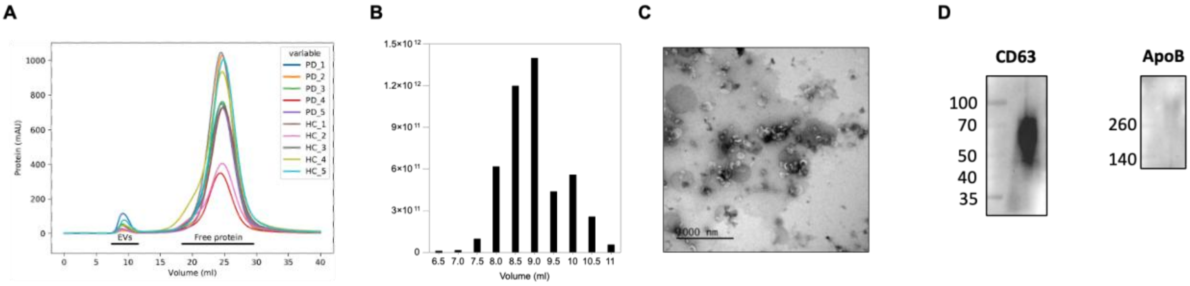


**Supplementary Figure 4:** Raw data of the αSyn SAA experiment with stool-derived EVs from Parkinson´s patients (A) and healthy controls (B). Each experiment was performed in triplicates (n=3). The threshold, indicated by a dotted line, was set at 30,000 RFU.

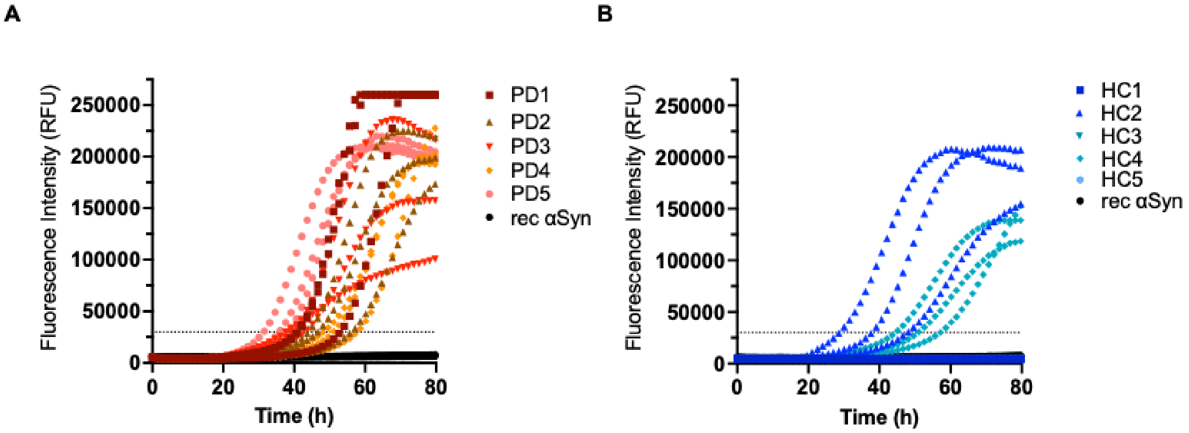

**Supplementary Figure 5:** Raw data of αSyn SAA experiments with pre-incubated recombinant αSyn. A–E) Raw αSyn SAA data after recombinant monomeric αSyn was pre-incubated with stool-derived EVs from Parkinson's disease (PD) patients. Each sample was run in triplicates (n=3). The threshold indicated by a dotted line, was set at 30,000 RFU. F) Graphical representation of TTT for each PD sample at different time points during the pre-incubation period.

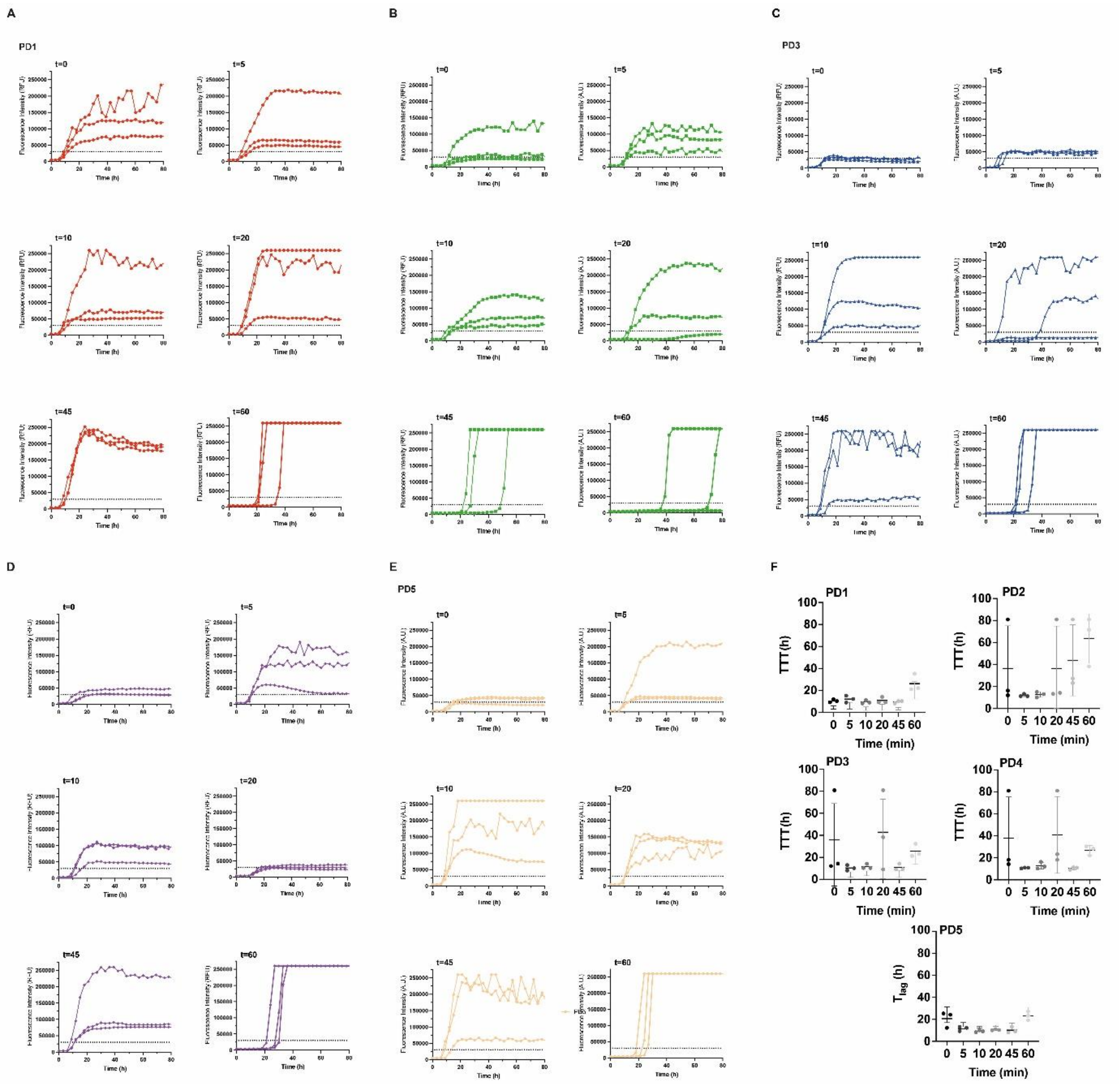

**Supplementary Figure 6:** Stool-isolated EVs from healthy controls (HC) were pre-incubated with recombinant monomeric αSyn for t=0 and t=60 min, followed by αSyn SAA (A). Graphs show TTT (Mann-Whitney U Test, ****$p$<0.0001) (B).

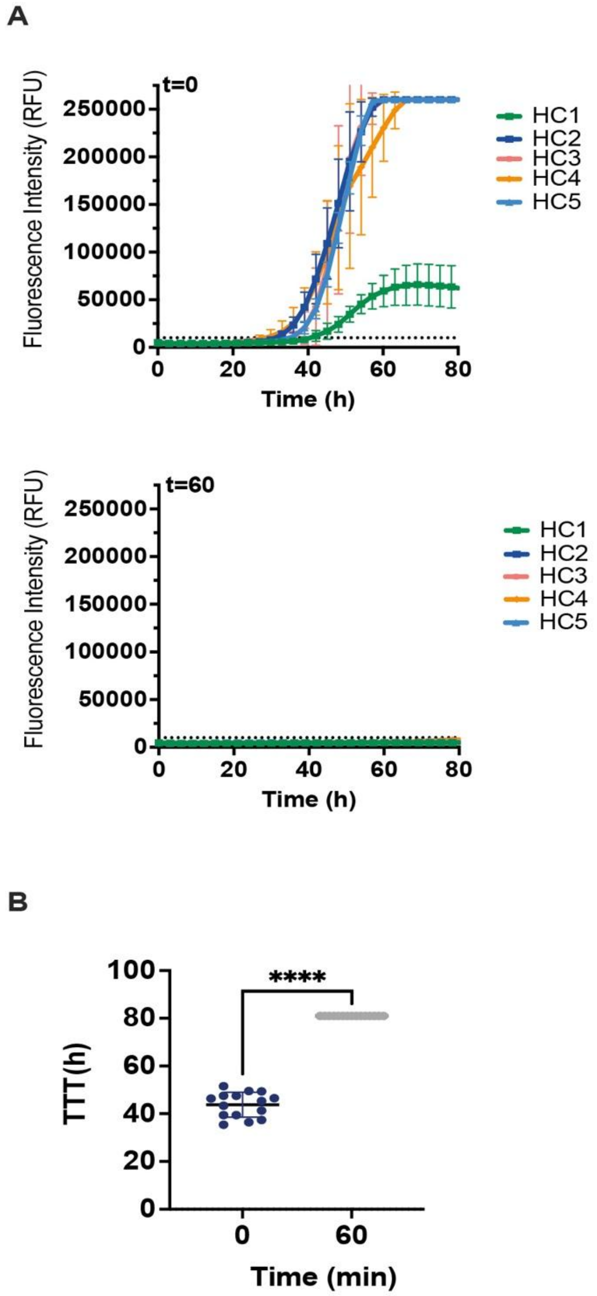
